# Seeing the forest and the trees: a workflow for automatic acquisition of ultra-high resolution drone photos of tropical forest canopies to support botanical and ecological studies

**DOI:** 10.1101/2025.09.02.673753

**Authors:** Etienne Laliberté, Antoine Caron-Guay, Vincent Le Falher, Guillaume Tougas, Helene C. Muller-Landau, Gonzalo Rivas-Torres, Thomas R. Walla, Hugo Baudchon, Mélvin Hernandez, Adrian Buenaño, Anna Weber, Jeffrey Q. Chambers, Jomber Chota Inuma, Fernando Araúz, Jorge Valdes, Andrés Hernández, David Brassfield, Paulo Araujo Filho, Vicente Vasquez, Adriana Simonetti, Daniel Magnabosco Marra, Caroline de Moura Vasconcelos, Jarol Fernando Vaca, Geovanny Rivadeneyra, José Illanes, Luis A. Salagaje-Muela, Jefferson Gualinga

## Abstract

Tropical forest canopies contain many tree and liana species, and foliar and reproductive characteristics useful for taxonomic identification are often difficult to see from the forest floor. As such, taxonomic identification often becomes a bottleneck in tropical forest inventories. Here we present a drone-based workflow to automatically acquire large volumes of close-up, ultra-high resolution photos of selected tree crowns (or specific locations over the canopy) to support tropical botanical and ecological studies (https://youtu.be/80goMEifpc4). Our workflow is built around the small, easy-to-use DJI Mavic 3 Enterprise (M3E) drone, which is equipped with a wide-angle and a telephoto camera. On day one, the pilot maps a forest area of up to ∼200 ha with the wide-angle camera to generate a high-resolution digital surface model (DSM) and orthomosaic using structure-from-motion (SfM) photogrammetry. On subsequent days, the pilot acquires close-up photos with the telephoto camera from up to 300 selected canopy trees per day. These close-up photos are acquired from 6 m above the canopy and contain a high level of visual detail that allows botanists to reliably identify many tree and liana species. The photos are geolocated with survey-grade accuracy using RTK GNSS, thus facilitating spatial co-registration with other data sources, including the photogrammetry products. The primary operational challenge of our workflow is the need to maintain RTK corrections with the drone to ensure that close-up photos are acquired exactly at the predefined locations. The maximum operational range we achieved was 3 km, which would allow the pilot to reach any tree within a ∼2800 ha area from the take-off point. Although our workflow was developed to support taxonomic identification of tropical trees and lianas, it could be extended to any other forest or vegetation type to support botanical, phenological, and ecological studies. We provide **`harpia`**, an open-source Python library to program these automatic close-up photo missions with the M3E drone (https://github.com/traitlab/harpia).

**Data/code for peer review statement:** We provide **`harpia`**, an open-source Python library to program these automatic close-up photo missions (https://github.com/traitlab/harpia). Drone imagery and labelled close-up photo data are not yet publicly available because they were acquired with the goal of publishing benchmark machine learning datasets and models for tree and liana species classification and prior publication of the data would jeopardize this future publication.

## Introduction

Tropical forests hold much of the Earth’s tree biodiversity (Cardoso et al., 2017; Gatti et al., 2022) and aboveground carbon (Feng et al., 2024; Santoro & Cartus, 2023). In particular, large canopy and emergent trees contribute disproportionately to tropical forest biomass, productivity, and functioning (Araujo et al., 2020; Lutz et al., 2018). Acquiring abundant species-specific data on tropical tree abundance, phenology, growth and mortality is critically important to predict ecosystem responses (Davies et al., 2021), but conducting tropical tree surveys is logistically challenging (ForestPlots.net et al., 2021). The exceptionally high diversity of tropical trees requires a high level of taxonomic expertise, and few people possess the necessary identification skills (Phillips, 2023). Complicating things further is the fact that the leaves, flowers or fruits that can assist tree identification, especially for large trees and lianas, are often high in the canopy, and cannot easily be collected or even viewed. For these reasons, taxonomic identification tends to be a major bottleneck in tropical tree surveys (ForestPlots.net et al., 2021).

Because of these difficulties, there has been much interest in developing remote sensing methods to identify tropical tree species from the air or even from space. Unfortunately, the relatively coarse spatial and/or spectral resolution of satellite imagery is rarely high enough to reliably identify individual trees to species (but see Ferreira et al., 2021; Kellner & Burley, 2024; Sánchez-Azofeifa et al., 2011). Airborne imaging spectroscopy has shown promise for tropical tree species mapping due its high spectral resolution (Féret & Asner, 2013), but this technology is very expensive and inaccessible to most researchers. In general, development of tools for taxonomic identification from airborne or spaceborne remote sensing requires high quality ground-truthing datasets, e.g., to build large spectral libraries linking crown reflectance to tree species identity (Asner & Martin, 2016).

Small unoccupied aerial vehicles (UAVs; hereafter, “drones”) are increasingly being used to survey trees and forests. These drones are now relatively inexpensive and easy to use. The most affordable drones are equipped with red-green-blue (RGB) cameras, which have low spectral resolution (only three bands in the visible range), but can offer very high spatial resolution. Such drones can be used to accurately map forest canopies (Fig. 1a-b) and acquire ultra-high resolution close-up photos of individual trees (Fig. 1c). Drone photos can be very useful to locate trees of conservation interest (Nyberg et al., 2024), monitor foliar (Park et al., 2019) or reproductive (Lee et al., 2023) phenology, estimate liana infestation (Waite et al., 2019), and quantify tree mortality (Mosig et al., 2024), among other use cases.

**Figure 1.**
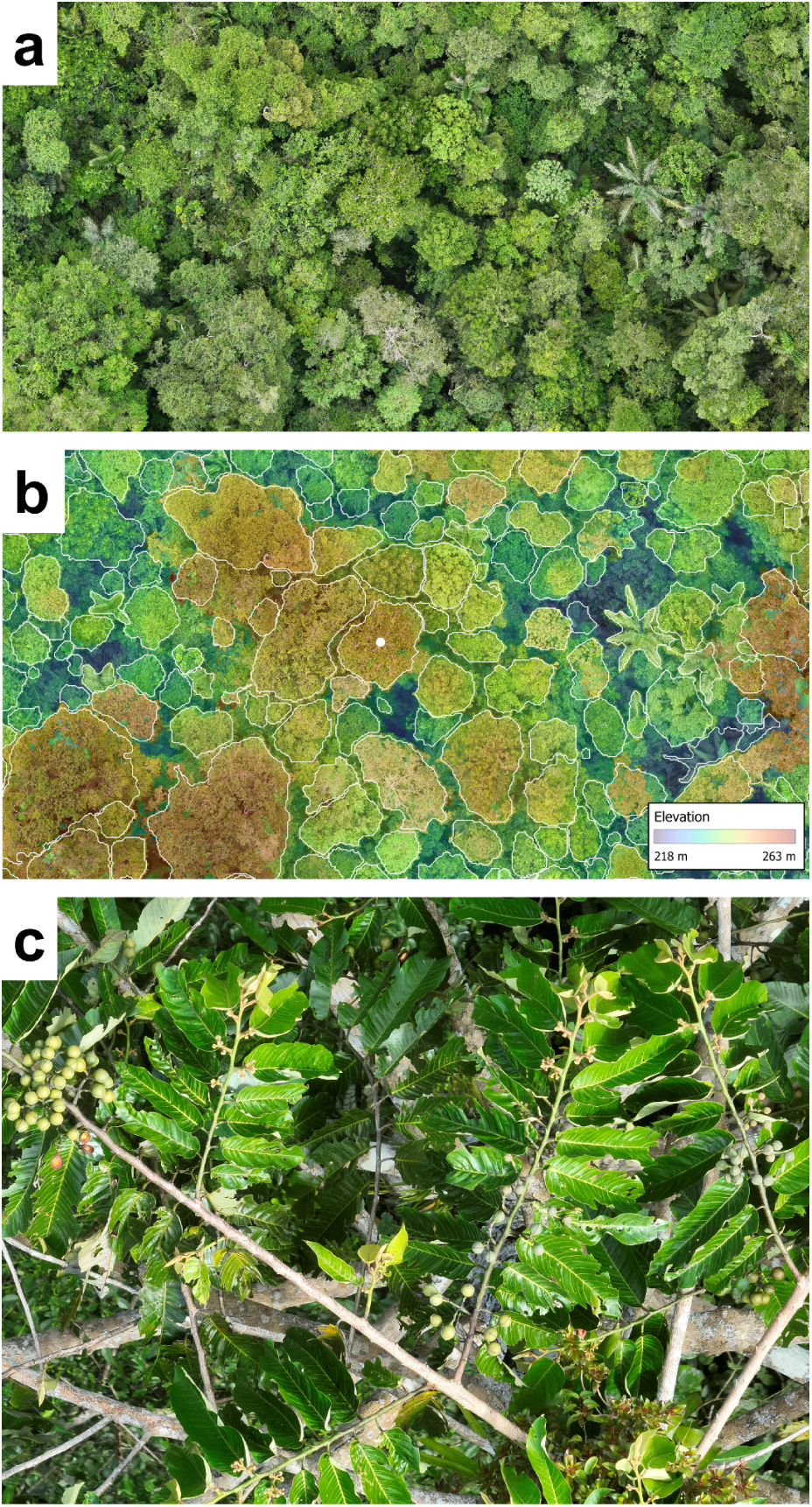
High-resolution RGB images of tropical forest canopies using drones at two different spatial scales at Tiputini Biodiversity Station, Ecuador. **a**) Subset of RGB orthomosaic (1.9 cm pixel^-1^) **b**) overlaid with crown segmentation polygons in white and the DSM at 30% transparency. The white dot shows the location of the close-up photo, in the center of a selected tree crown. **c**) Close-up photo acquired with the DJI M3E telephoto camera, 6 m directly above the tree shown with a yellow dot in **a**). The tree could be identified as *Virola obovata* Ducke from the close-up photo, as foliar characteristics and fruits are clearly visible. Photos credits: **a**) and **b**) José Illanes, **c**) Jefferson Gualinga.

It is fairly easy to fly a drone manually toward selected trees and take a close-up photo of their crowns. Such a manual approach can work well when the number of trees to survey is small and when they are close to the take-off point, so that the pilot can maintain strong video signal between drone and remote controller (Nyberg et al., 2024). However, for larger-scale tree surveys over broader areas (i.e. tens or hundreds of hectares), or when the number of trees to be studied lies in the hundreds or the thousands, this manual photo acquisition approach becomes inefficient at best or infeasible at worst. In these situations, using an automated, accurate, repeatable photo acquisition workflow becomes highly desirable.

We present a custom workflow for the automatic acquisition of close-up drone photos of selected tree crowns (or specific locations in the canopy) to support tropical botanical and ecological work. Our workflow requires a prior drone mapping mission to be completed over the area of interest, from which a digital surface model (DSM) and an RGB orthomosaic is produced, to guide the acquisition of close-up photos. Our workflow then allows one pilot to photograph the crowns of up to 300 selected canopy trees per day within this area. These close-up photos are acquired from 6 m above the canopy and contain a high level of visual detail that allows expert botanists to reliably identify many tree and liana species (Fig. 1c). Our workflow was initially developed as part of the winning team of the XPRIZE Rainforest competition (https://www.xprize.org/prizes/rainforest) to support taxonomic identification of tropical canopy trees, and was then further refined after the competition. We have used it successfully in over 150 missions in three tropical countries (Panama, Brazil, Ecuador) to acquire photos from over 15,000 trees. Although our workflow was initially designed for tropical forests and trees, it could be applied to any other forest or vegetation type to support floristic, phenological, or ecological studies.

Close-up canopy photos can also support the study of lianas and (hemi)epiphytes. We provide the **`harpia`** open-source Python library (https://github.com/traitlab/harpia) to generate automatic close-up photo missions (i.e. automatic flight plans) that can be imported in the drone remote control software. The name **`harpia`** comes from the neotropical harpy eagle (*Harpia harpyja*), which like our drone has very high visual acuity to observe the canopy with great detail.

## Materials and Methods

### Equipment

Our workflow is based around the DJI Mavic 3 Enterprise (M3E) drone, which is equipped with a dual wide-angle and a telephoto camera (DJI, China, https://enterprise.dji.com/mavic-3-enterprise). The M3E drone is small, portable, relatively easy to use, and widely accessible due to its relative affordability (for an enterprise drone) and global availability. Drone technology evolves rapidly, and we expect that our workflow can be adapted to other similar drones with comparable cameras and mission planning capabilities (e.g., the DJI Matrice 4 Enterprise that recently came out; https://enterprise.dji.com/matrice-4-series). Here we focus exclusively on the M3E, which we tested extensively in tropical and temperate forests since 2023.

Our workflow relies on the real-time kinematic (RTK) GNSS positioning ability of the M3E to send the drone back to accurate locations for the acquisition of close-up photos. We equip the M3E with the Mavic 3 Enterprise RTK Series module. We also use the DJI D-RTK2 GNSS base station (https://www.dji.com/ca/d-rtk-2) to send RTK corrections to the drone, although it is also possible to use another GNSS base station that can broadcast local NTRIP corrections, such as the Emlid RS2/RS2+ receiver (https://docs.emlid.com/reachrs2/integration/dji-rtk). In locations with internet access, we have also been able to use NTRIP RTK corrections broadcasted over the internet when those were available. We use the DJI RC Pro remote controller that comes with the M3E, running the most recent version of DJI Pilot 2 (v. 12.15.0.112) at the time of writing.

Extensive prior experience in drone operation is not required to implement our proposed workflow. We estimate that, with appropriate training on the use of the M3E drone, relevant local laws and permitting procedures, and approximately three to four weeks of practical instruction focused on takeoff, landing, in-flight control, and emergency response, operators should be fully capable of capturing the initial imagery required for this workflow.

### Mapping

Planning close-up photo missions within an area of interest first requires a high-resolution and accurate DSM from that area. The DSM is needed to extract the ellipsoidal elevations to use at selected locations where close-up photos are to be acquired. A co-aligned high-resolution RGB orthomosaic can also be useful when selecting these locations to provide forest canopy context. In practice, the DSM (and the RGB orthomosaic) will most often come from a previous mapping mission done with the same M3E drone (using its wide-angle camera). However, it could come from other sources (e.g., a previous LiDAR mission), as long as the DSM is accurately georeferenced and reflects the current state of the canopy. It is important that the DSM represents the forest canopy surface as accurately as possible at the time of the acquisition of the close-up photos, because the close-up photos are programmed to be acquired at only 6 m above the canopy, and changes in canopy structure could increase the risk of drone collisions or lead to more distant photos that provide less detail.

If the mapping mission is done with the M3E, we recommend the following acquisition key parameters in DJI Pilot 2:

- Front overlap: ≥85%
- Side overlap: ≥75% (high side overlap helps with photogrammetric reconstruction but increases flight time)
- Flight altitude: ≥50 m above canopy (lower generates more visual artefacts, too high leads to lower ground sampling distance or GSD)
- Altitude reference mode: Real-time terrain follow (helps to maintain constant overlap and GSD)
- Shutter speed: priority, at 1/1000 s (or faster if sunny)
- RTK: ON, and ensure that M3E position is FIX (alternatively, use PPK later; https://enterprise-insights.dji.com/blog/ppk-post-processed-kinematics-workflow)

### Photogrammetry

Once photos from the mapping mission are acquired, any structure-from-motion (SfM) photogrammetry software could be used (Li & McKinney, 2022). We use Agisoft Metashape, which in our experience gives good results for forest canopies, and is commonly used in forest studies (e.g., Cloutier et al., 2024; Schiefer et al., 2020; Tinkham & Swayze, 2021). A good reference for the optimal Metashape parameters to choose to ensure a good reconstruction of trees is Tinkham and Swayze (2021).

The two primary considerations for the photogrammetry processing are: (1) to obtain a high-quality DSM in which the individual trees and canopy are reconstructed as faithfully and accurately as possible, with as few gaps or visual artefacts as possible, and (2) to ensure that the DSM is georeferenced accurately across the entire area of interest. Meeting this last criterion is facilitated by the proprietary TimeSync 2.0 technology of the DJI M3E, which synchronizes data from the GNSS antenna, gimbal, camera and IMU at the microsecond level to ensure that the geotag of each photo is in the optical center of each image and survey-grade accurate. The TimeSync technology essentially removes the need for ground control points (Kalacska et al., 2020), which are very difficult or even impossible to position in tall, dense tropical forests. Importantly, this helps to ensure 3D alignment between the DSM and the selected locations for the close-up photos.

### Selection of trees for close-up photos

Once the DSM (and the RGB orthomosaic if available) is produced, it can be loaded in GIS software (e.g., QGIS; https://qgis.org). The user creates a point layer and creates one point for each selected location where a close-up photo is to be acquired. If individual crowns are indicated by polygons (e.g., tree bounding boxes or crown segmentation polygons, as in Fig. 1b), then their centroids can be used as horizontal coordinates for the planned photos. Recent artificial intelligence (AI) models can help to provide such automatic tree crown bounding boxes or segmentations from high-resolution RGB orthomosaics of tropical forests (Baudchon et al., 2025), and we used these models to guide tree selection in the present study.

The selection of trees to be mapped can follow either taxonomic criteria (e.g., individuals from the same genus with similar crown architecture) or ecological traits (e.g., emergent trees, or crowns with conspicuous flowers or fruits). Sometimes tree crowns show gaps in foliage, and it can be useful to leverage the RGB orthomosaic to select specific locations within the crown that have sufficient leaf cover to facilitate species identification. Additionally, if tree crowns are visibly covered by lianas, it may be useful to select locations of close-up photos away from those lianas if the goal is to identify the host tree (or vice-versa if the goal is to study lianas).

Selection of locations for close-up photos should also consider spatial and structural attributes of their location, especially in relation to neighbouring trees and topographical position. For instance, trees with heavily interwoven crowns, those with protruding branches extending beyond the main canopy, or individuals located in valleys or in close proximity to taller neighboring trees should be avoided for drone safety considerations. In addition, the density of the forest and variations in topography may block RTK signals, potentially resulting in signal loss or disconnection. These issues are discussed further in the Discussion section.

### Close-up photo missions

#### Software

Our Python library, **`harpia`** (https://github.com/traitlab/harpia), requires a point (or polygon; in which case the polygon representative point will be used as the points) layer and a DSM (stored as a single-band GeoTiff). Our algorithm will generate a `.kmz` waypoint mission file that can be imported in DJI Pilot 2 on the remote controller of the M3E. Our algorithm extracts the maximum elevation within a 3 m circular buffer around each selected point from the DSM, and then adds 6 m. It uses those maximum elevations + 6 m, along with their horizontal coordinates, as the 3D locations where the drone will acquire close-up photos.

To determine the shortest, most efficient flight path between the selected photo locations, we use the Traveling Salesman Problem (TSP) algorithm. The starting point can be decided by the user. Additional waypoints are added as intermediate locations as the drone will travel from tree to tree in order to ensure a safe travel path (Fig. 2). Once the `.kmz` is imported in DJI Pilot 2, the user can run this waypoint mission automatically.

**Figure 2.**
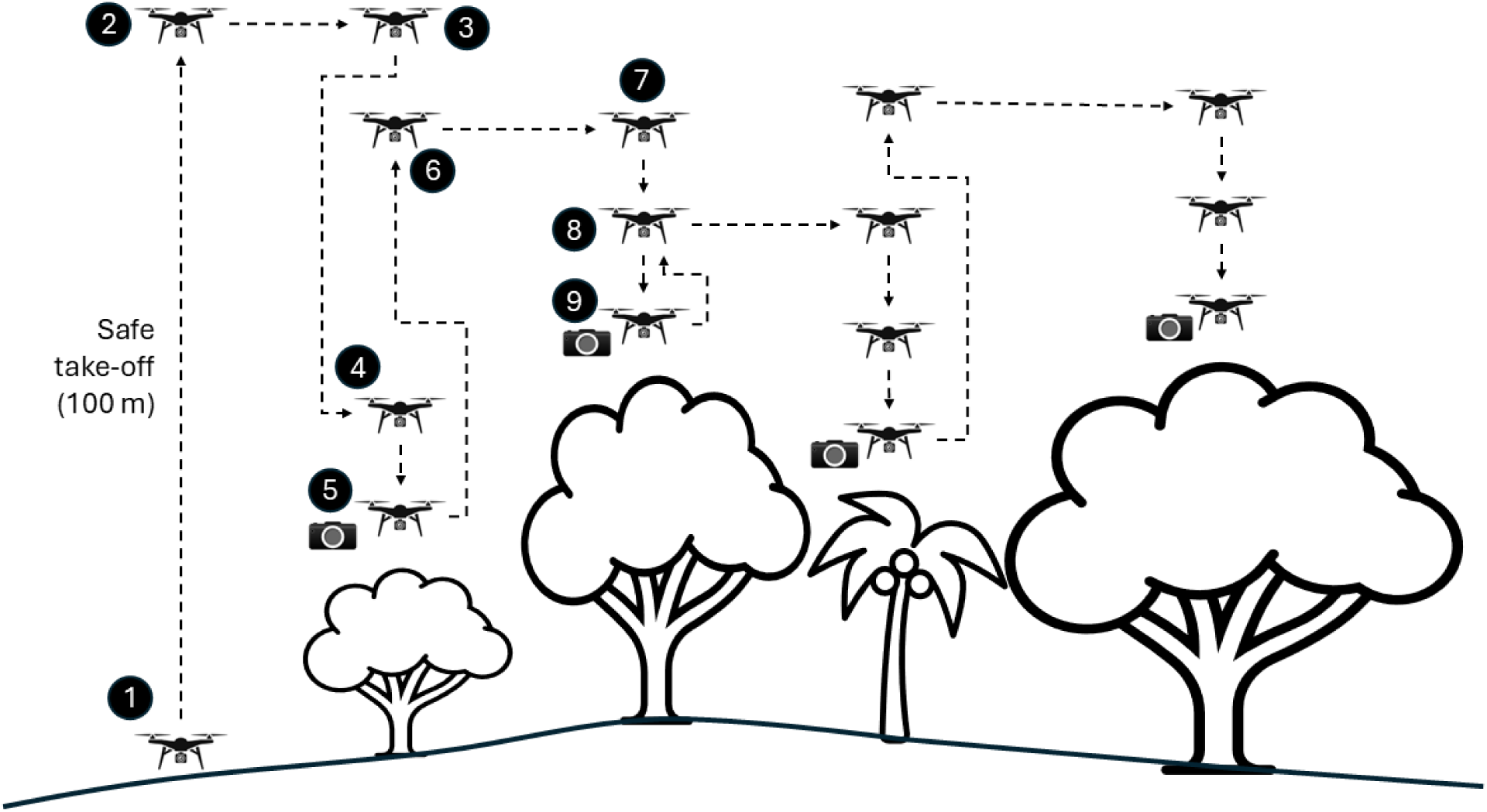
Diagram showing an automatic close-up photo mission. (1) The drone takes off from an open area and (2) flies to the safe take-off altitude of 100 m. (3) It moves at that altitude to the first selected tree, and then (4) descends at maximum speed to 10 m above the selected photo location. (5) The drone then descends at a slower speed to the selected photo location, which is 6 m above the maximum elevation of the canopy in a 3-m circular buffer around the selected location, and takes the photos. (6) The drone then flies straight up to a safe altitude that is 16 m above the highest elevation in a 10-m wide straight corridor between the first and (7) second tree. (8) Again, it descends at maximum speed to the approach point, and then descends to (9) 6 m above the second tree to take photos. Because the third tree is lower than the second and there are no higher obstacles in between, it flies straight up back to (8) before travelling to the third tree. The same logic applies to the fourth, and all subsequent trees in the mission. A video showing this workflow in action can be found here: https://youtu.be/80goMEifpc4.

#### Mission description

The automatic close-up photo mission can be summarized as follows (Fig. 2). First, the drone climbs vertically to the default safe take-off altitude of 100 m (this can be changed by the user) above the take-off point. Then, the drone will move to the first waypoint, which is directly above the first selected tree (or point). The drone will point its camera down -90°, and will then descend at its maximum speed of 6 m s^-1^ to 10 m above the target location. It will then move to the target location (i.e. the position where the drone takes the photo, which is 6 m above the maximum elevation within a 3-m circular buffer around the selected location of the canopy) at a slower approach speed of 3 m s^-1^. Before taking photos, the drone reorients its heading to face geographic north (0°). This is done so that the close-up photos are taken with consistent orientation and can easily be aligned to the orthomosaic.

Once oriented to geographic north, the drone then takes two photos: first, a photo at maximum optical zoom (using the telephoto camera), and then a photo at minimum zoom to provide context (using the wide-angle camera). The drone will then ascend back up to a safe elevation which is determined as the maximum elevation within a 10-m corridor between that tree and the next one to photograph (Fig. 2). As the drone moves between trees, the camera is set to the wide angle camera and the gimbal angle is set at -15° to provide a good view of the canopy and any potential obstacles to the pilot. A video showing an example mission at the Tiputini Biodiversity Station in Ecuador can be viewed here: https://youtu.be/80goMEifpc4.

#### RTK corrections

Prior to launching the drone for the close-up mission, the pilot needs to ensure that the M3E receives RTK corrections and that its position status is FIX. RTK corrections need to be received at all times during the close-up photo mission to ensure position accuracy. RTK FIX can be maintained for up to 10 min after signal loss if “Maintain Position Accuracy” is activated in DJI Pilot 2. If the take-off point is located within the area of the DSM, we strongly recommend that the pilot confirms that the elevation of the M3E at take-off point matches the expected elevation from the same point on the DSM, to ensure spatial alignment.

#### Spatial reference system

The spatial reference system of the RTK base station needs to be the same as that of the DSM used to plan the close-up photo mission. Importantly, **`harpia`** requires WGS84 (or ITRF2020, which is considered equivalent to WGS84) *ellipsoidal* (and *not* orthometric) elevations for both the DSM (mapping mission) and for the close-up photo elevations. If the coordinates of the base station used for the mapping mission were not determined with absolute accuracy, then relative accuracy needs to be set by positioning the base station in the exact same location that was used for the mapping mission. In this case, the exact same base station coordinates as used for the mapping mission need to be entered in DJI Pilot 2. This helps to ensure that every close-up photo is accurately positioned *relative* to the base station and/or the DSM.

Failure to ensure accurate spatial alignment between the DSM and the close-up photo locations will result at best in photos being taken from different locations than those intended, and at worst could result in the drone colliding with a neighbouring tree (but see *Drone safety considerations* in the Discussion). If a ground survey marker with known coordinates is used to position the DJI-RTK2 base station on, it is important to note that DJI Pilot 2 uses as base coordinates the GNSS *antenna* elevation, not the *ground* elevation. Since the DJI-RTK2 antenna is located 1.8 m above the ground when positioned with its pole and tripod, then 1.8 m must be added to the known ground elevation in DJI Pilot 2.

#### Camera parameters

It is important to obtain the highest possible image quality for close-up photos to facilitate taxonomic identification. We typically acquire close-up photos with shutter speed priority set around 1/160 s (in overcast conditions with diffuse light, which are best for image quality). This helps to maintain the ISO around 100, which is ideal to minimize digital noise in the photos. Because the drone is hovering as it is taking the close-up photos, any motion blur will be due to the wind and not the movement of the drone itself, such that a slower shutter speed can be used than for a mapping mission. We have not found motion of foliage from the drone propellers to be an issue. We also recommend setting the focus mode of the telephoto camera to continuous auto-focus (AFC) prior to starting the mission to help ensure that foliage remains in focus as the photo is taken.

### Taxonomic identifications of photos

We used the Labelbox (https://labelbox.com) platform for expert botanists to provide taxonomic identifications of trees and lianas from the close-up photos. Labelbox is free for research and education use (https://labelbox.com/research). We set up image labelling projects using instance segmentation labelling tools. We added nested classifications to the segmentation tools with taxonomic lists provided by the experts. These lists included scientific names for species, genera and families that are recorded in the region. Botanists drew the segmentation masks for trees or lianas and added identifications to the taxonomic level that they could reliably identify. Images were labelled by one expert botanist familiar with the local flora, and identifications and segmentation masks were then reviewed by at least one additional person.

### Species accumulation curves

In order to explore how many tree and liana species are captured per sampling effort (i.e. number of close-up photos taken), we generated species accumulation curves for the two sites where we acquired and identified a large number of close-up photos: Barro Colorado Island (Panama) and Tiputini Biodiversity Station (Ecuador). We considered a subset of 2207 close-up photos from both of these sites in which at least one tree or liana taxon was identified to species. Species accumulation curves were created using the `specaccum` function of the `vegan` package (Oksanen et al., 2025).

## Results

Using our **`harpia`** workflow, we were able to acquire close-up photos from more than 15,000 individual trees using six different M3E drones at tropical sites in three countries (Panama, Ecuador, Brazil). This corresponded to a total of 153 individual drone missions (one mission can use several batteries). Examples of close-up photos are shown in Figure 3.

**Figure 3.**
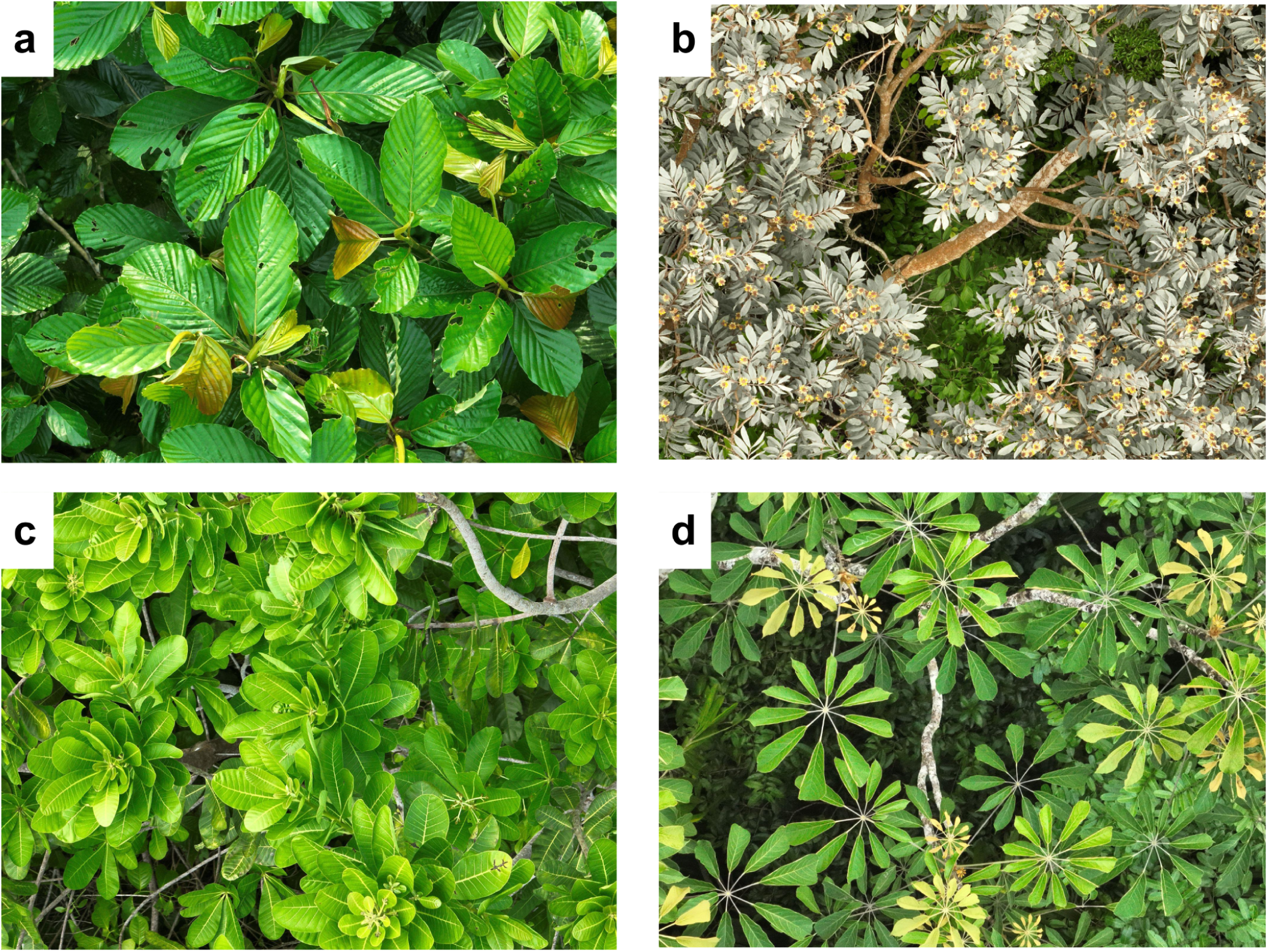
Examples of close-up photos from different tree species and sites **a**) *Pourouma bicolor*, Tiputini Biodiversity Station, Ecuador. **b**) *Tachigali myrmecophila*, ZF-2 station, Brazil. **c**) *Anacardium excelsum*, Barro Colorado Island, Panama. **d**) *Didymopanax morototoni*, Finca Roubik 1-ha plot, Panama. Photos credits: **a**) Adrian Buenaño, **b**) Anna Weber, **c**) Mélvin Hernandez, **d**) Vicente Vasquez.

Out of these >15,000 close-up photos of trees (Tables 1 and 2), to date a total of 2524 have been labelled by expert botanists (Table 1). 80.39% of these labelled images showed a single tree species, whereas 19.61% showed two or more species, most often including one or more liana (i.e. woody vine) species. Of the 401 distinct taxa that could be identified in the images, 86.70% of the labelled images were identified at species level, 9.92% at genus level, and 3.38% at family level (Table 1). At the species-rich Tiputini Biodiversity Station in Ecuador, some trees that could only be identified to genus are probably undescribed species.

**Table 1.** Statistics of drone close-up photos at the two sites where photos received the largest labelling effort by local botanists, as of September 2, 2025.

| Site | Barro Colorado Island | Tiputini Biodiversity Station |
| --- | --- | --- |
| Number of close-up photos taken | 4485 | 6531 |
| Number of close-up photos labeled | 1763 | 1436 |
| Photos labeled with more than one taxon (%) | 22.86 | 8.64 |
| Labels identified to species (%) | 91.10 | 76.73 |
| Labels identified to genus (%) | 4.94 | 18.63 |
| Labels identified to family (%) | 3.96 | 4.64 |
| Number of distinct species | 168 | 218 |
| Number of distinct genera | 121 | 173 |
| Number of distinct families | 48 | 62 |

**Table 2.** Number of drone close-up photos acquired for sites that have not yet been labelled by botanists.

| Site | Number of close-up photos |
| --- | --- |
| AmazonFACE plots, Brazil | 219 |
| EEST, Brazil | 884 |
| Adolpho Ducke Forest Reserve, Brazil | 189 |
| Various sites in Central Panama | 726 |
| Yasuní 25-ha plot, Ecuador | 3225 |

No species labels were supplied for about 4% of photos that were viewed by botanists. This includes cases in which the photo was not properly exposed (generally over-exposed), out of focus or too low resolution (e.g., due to the drone flying too high). This also includes cases in which there were many intermixed species and the photo was considered too complicated to be worth the botanist’s time, and cases in which the visible plants could not be identified at least to family level by the botanist. Failure to identify taxa could be due to dead or leafless trees lacking distinguishing features of their bark, cases in which multiple species from different families have leaves that cannot be distinguished at this image resolution, as well as cases of rare species not known to the botanists.

Species accumulation curves showed the expected rapid initial increase in number of tree and liana species with increasing sampling effort, which then tapers off (Fig. 4). The total number of species and the rate at which new species were found were considerably higher at the Tiputini Biodiversity Station than at Barro Colorado Island in Panama (Fig. 4). These results are consistent with the exceptionally high tree species diversity known to occur in the Yasuní region in which Tiputini Biodiversity Station is located (Valencia et al., 1994).

**Figure 4.**
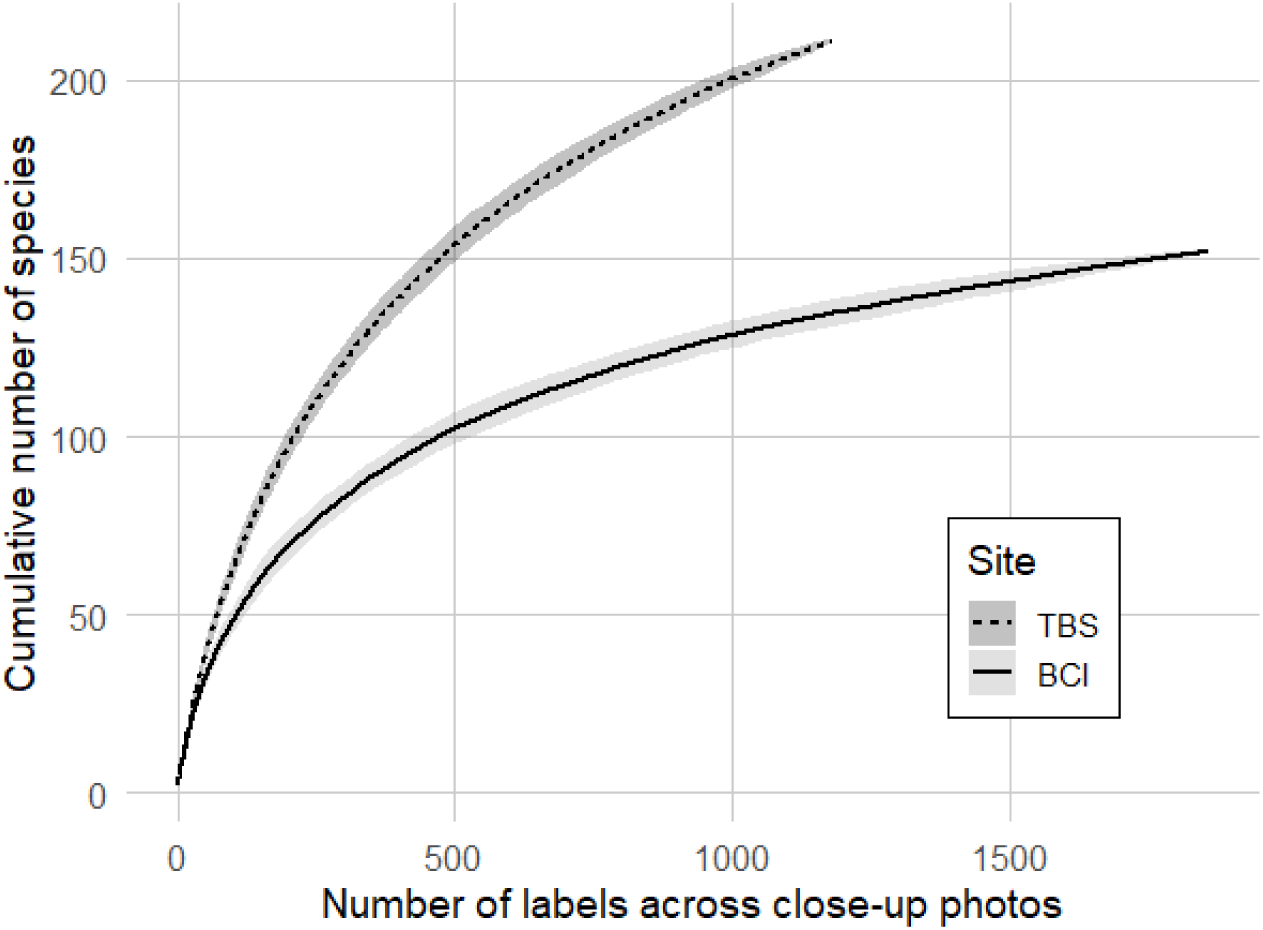
Species accumulation curves at Tiputini Biodiversity Station (TBS), Ecuador and at Barro Colorado Island (BCI), Panama. Shaded areas represent the standard deviation of estimates.

The geographical coordinates of each close-up photo stored in the EXIF metadata were very accurate (Fig. 5). When comparing the distance between the coordinates of each photo acquired in the EXIF metadata and the intended location selected to send the drone, the absolute distance was generally less than 70 cm horizontal and 60 cm vertical. The mean horizontal error was 12.5 ± 7.5 cm and the mean vertical error was 4.7 ± 5.5 cm (mean ± SD).

**Figure 5.**
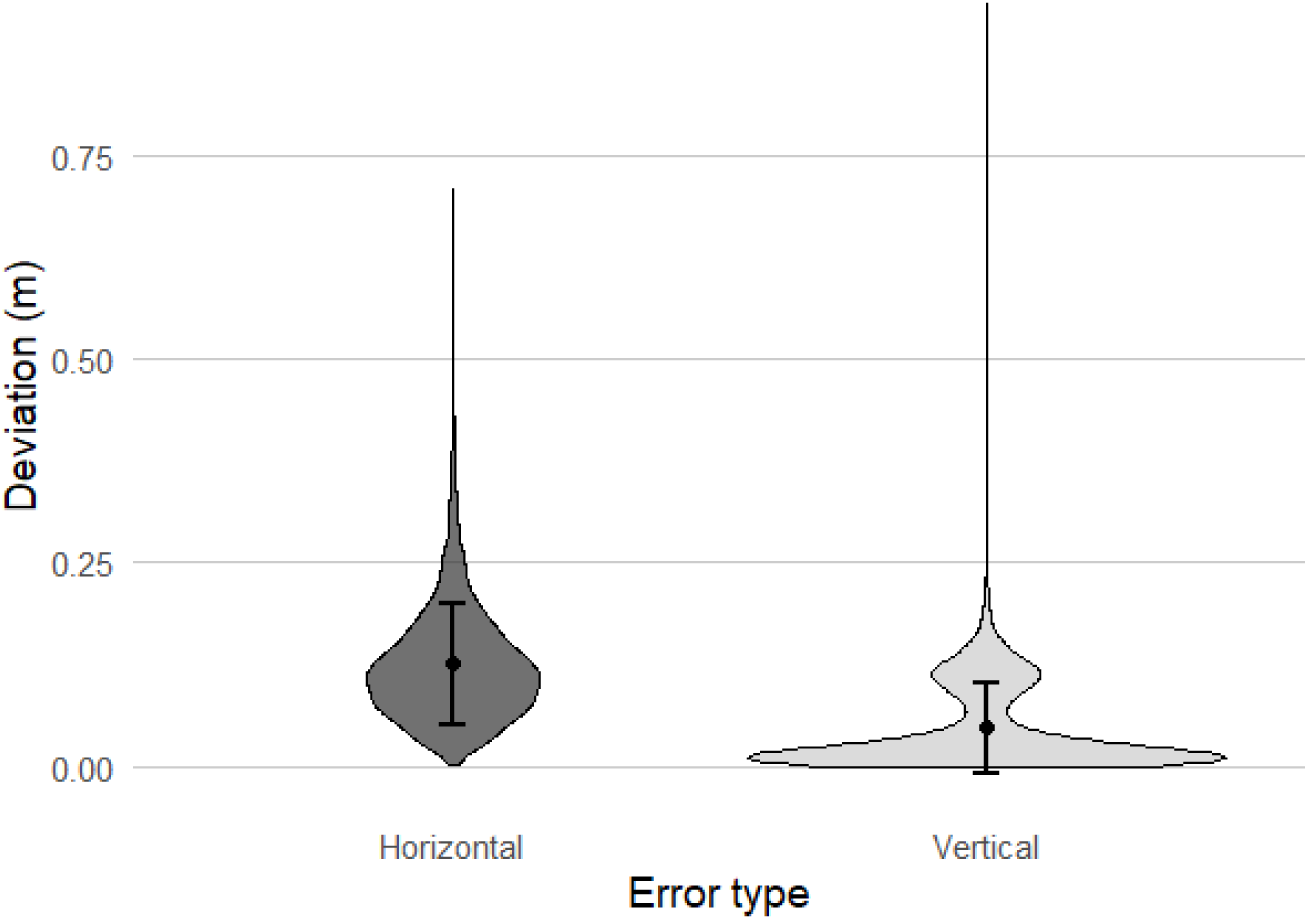
Absolute distance (i.e. deviation or location error) between specified coordinates and actual drone position for a subset of 3812 images that were captured with the most recent version of **`harpia`**.

We were able to acquire close-up photos from a maximum of 42 trees using a single M3E battery, which corresponds to approximately 30 min of flight. However, when the distance between take-off and selected trees was large, this number could be as low as 15 trees per battery. It is recommended to return with >20% battery for drone safety. In practice, our M3E kits often contain seven batteries each as this is the maximum number of batteries that fit in the M3E case. If the pilot is able to charge batteries during the day (e.g., using a portable charger), then it is possible to use ∼10 batteries or so in one day, which gives a total of 5-6 h of flight time. The maximum number of trees we have photographed by one pilot in one day is 300. To achieve this, flights were performed from 9:46 to 12:31, then from 16:04 to 17:32, corresponding to 4.22 hours of flight time.

The physical operational range of our close-up photo workflow is primarily limited by the RTK signal that needs to be maintained between the drone, the remote controller, and the RTK base station. In some jurisdictions, operational range may be further limited by legal constraints about maintaining the drone in visual-line-of-sight (VLOS). When present, towers can be used to solve this problem. For example, at Tiputini, we used a canopy observation platform as the launching point (Fig. 6a) to maximize signal strength between the drone, pilot and base station, and with this setup we were able to acquire close-up photos of canopy trees up to ∼3 km away. This gives an area of operation of ∼2800 ha around the take-off point.

**Figure 6.**
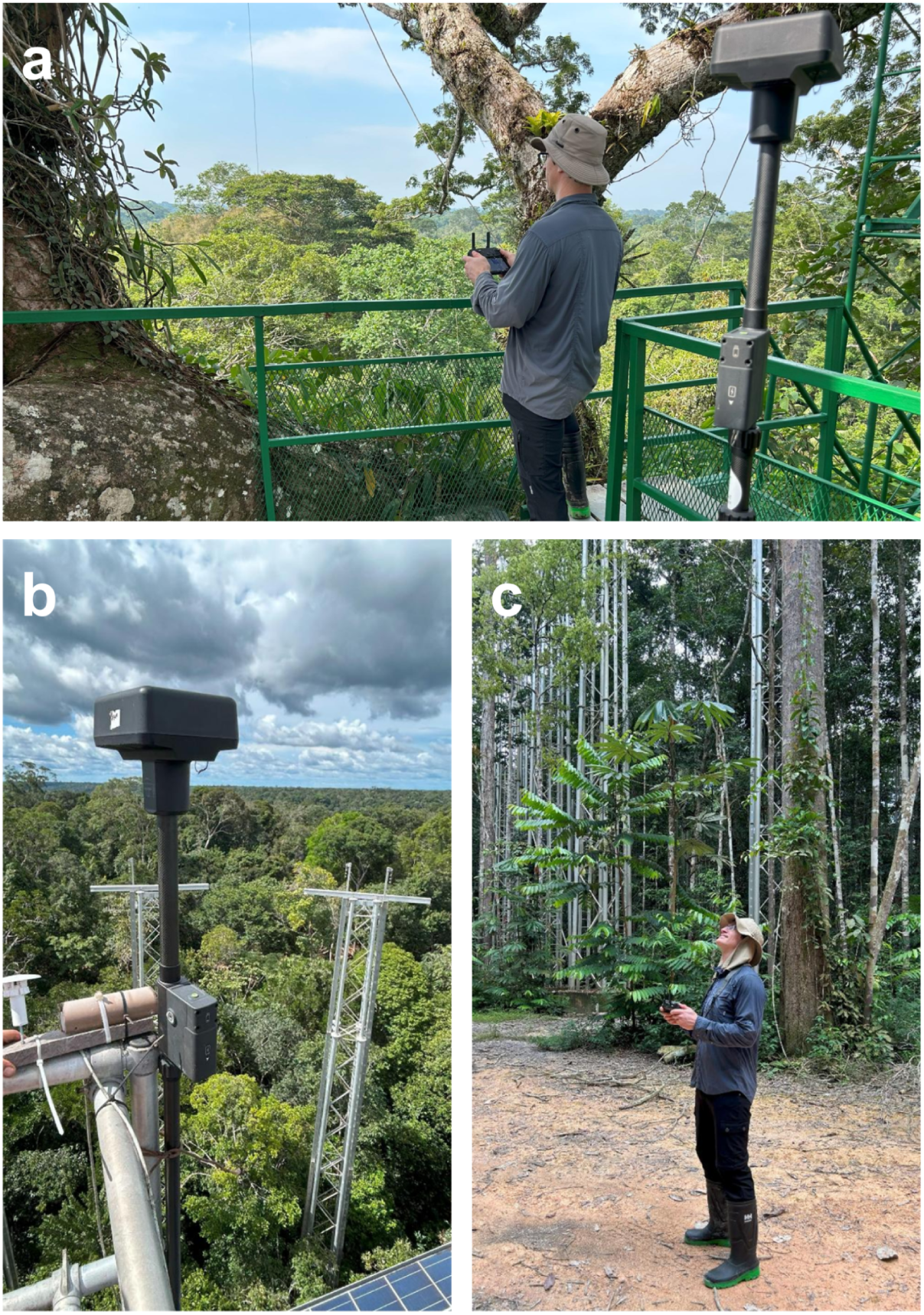
Using towers to improve communication between drone, remote controller and base station. (**a**) At the Tiputini Biodiversity Station, very few open areas on the ground can be used to install the GNSS D-RTK2 base station to view enough satellites. In addition, loss of signal is an issue if the pilot is on the ground. The solution we used was to position both the GNSS base station and the pilot at the top of the canopy tower. This 45-m tower is physically anchored to a tall, emergent kapok tree (*Ceiba pentandra*) that gives a clear visual line-of-sight between the pilot and the drone over several km. Two challenges with this setup are: the need to manually fly the drone when taking off and landing to avoid the many obstacles (branches, platform structure), and the relatively poor GNSS signal due to signal obstruction by the kapok branches over the GNSS base station. (**b**) At the AmazonFACE site, we used a tower to position the base station so that it had a clear view of the sky, since not enough satellites could be seen from the ground from the few open areas we could access. (**c**) However, at AmazonFACE we could still position the pilot on the ground, since we could maintain good communication between the remote controller and the base station. This setup was more convenient for the pilot, but we had to fly the drone up above the canopy before turning RTK on and getting an RTK FIX, as it could not see enough satellites while on the ground.

## Discussion

We presented a drone-based workflow to automatically acquire close-up photos of tropical tree crowns (https://youtu.be/80goMEifpc4). Our workflow is based on the DJI M3E drone and we provide **`harpia`**, a Python library (https://github.com/traitlab/harpia) to program automatic close-up photo missions using this drone. Our workflow was tested extensively in tropical forests in Panama, Ecuador and Brazil, and in temperate forests in Canada. We have been able to acquire close-up photos from >15,000 canopy trees in these forests from 153 distinct missions. The majority of the close-up photos that botanists reviewed could be reliably identified to species, a minority only to genus level, and even fewer only to family. The close-up photos contain sufficient detail to allow different applications, such as phenological observations of fruits or flowers, or estimations of foliar damage (e.g., herbivory). Below we describe some limitations and practical considerations of our workflow.

### Limitations of RTK

The primary operational challenge of our workflow is the need for the drone to receive RTK corrections and maintain FIX status during the entire mission to ensure that close-up photos are acquired exactly at the predefined locations. First, this requires that the GNSS D-RTK2 base station that sends RTK corrections is positioned in an open area. Ideally, this open area would give the GNSS base station antenna a full 360° view of the sky above 15° degree elevation. However, such open areas can be difficult to find in a dense tropical forest setting. In practice, we have successfully used a range of large canopy gaps, clearings, or roads that did not present such ideal open conditions. However, this leads to fewer satellites than can be seen at any given time by the base station, which can slow down or even prevent RTK FIX status. Second, the GNSS D-RTK2 base station must maintain strong communication with the remote controller at all times. The claimed maximum interference-free distance between remote controller and the D-RTK2 station is 200 m (https://www.dji.com/ca/d-rtk-2/info#specs), but in a dense tropical forest setting, this distance can be significantly shorter (i.e. tens of meters).

During the mission, the drone should always have a good view of the sky since it is above the canopy. However, if taking off from a smaller canopy gap, sometimes the drone may not see enough satellites while it is on or near the ground because of canopy obstruction. In our experience, in these situations, turning off RTK and flying the drone manually above the canopy, then turning RTK back on while it is hovering, can lead to a much faster RTK FIX. The FIX status can persist for a few minutes even after the drone is brought back to the ground.

When positioning the base station and pilot in an open area surrounded by trees, and launching the drone from that open area, the range of operation will be limited by the video and RTK signal blockage from the vegetation (and/or any surrounding hills). In practice, we noticed significant signal loss at around 600 m from the take-off point when launching from a canopy gap or other open area surrounded by forest. The M3E can maintain its high-precision position for up to 10 min after the RTK signal is weakened or lost. In **`harpia`**, the user can therefore choose to have the drone climb up to a high location every *n* (e.g., ten) trees to regain RTK signal between the drone and the base station in case signal was lost during the flight; this can happen when the drone is moving down toward a tree.

When towers or other tall structures are available, these can be used to significantly improve the communication between the drone, remote controller and base station (Fig. 6), and increase the range of operation of our solution. The ∼3 km maximum range we obtained at Tiputini with the canopy tower (Fig. 6a) could probably be extended even further. Indeed, the claimed maximum signal transmission unobstructed range (without any interference) of the M3E is 8-15 km (https://enterprise.dji.com/mavic-3-enterprise/specs). However, in practice the operational range may be further limited by local regulations about maintaining visual line of sight (VLOS) flights. Another consideration is that when positioning the base station on a tower, the GNSS D-RTK2 base station needs to be stable at all times. If there is too much movement (e.g., due to wind, or people moving on the tower), then this can lead to positioning errors.

We have also successfully used the DJI Cellular Dongle on the M3E in one site where cellular signal was available. This setup allows one to maintain video signal and RTK corrections to the drone via the cellular network when the Occusync 3 transmission between remote controller and drone is lost. However, it is only possible for the M3E to receive NTRIP RTK corrections over the internet with this setup; D-RTK2 corrections cannot be used. Such NTRIP corrections over the internet can be obtained by a commercial provider, a government network, or by broadcasting your own NTRIP corrections (e.g., Emlid NTRIP Caster; https://emlid.com/ntrip-caster/). In theory, the physical operational range when using this setup is only limited by the battery (i.e. flight time) and the availability and strength of the cellular signal to the drone, and the internet connection to the remote controller.

### Linking close-up photos to selected trees or other geospatial data

It is often desirable to link the close-up photos to a specific tree from which other measurements have been made. For example, there is often interest in linking ground measurements (e.g., diameter at breast height) of trees from forest plots to close-up photos from these same trees. In our experience however, it can be very difficult to spatially co-register close-up photos to ground measurements, which are often georeferenced using standard GNSS receivers under a dense canopy, or via more traditional surveying approaches. The geolocation of the close-up photo in its EXIF metadata, if acquired with RTK FIX from a base station whose coordinates are measured with survey-grade accuracy, will be very accurate (e.g., only a few cm error). In contrast, GNSS measurements from under a canopy suffer from poor GNSS signal and multipath errors and can be many meters off or more. In addition, tree trunks are rarely perfectly straight, so the crown center from above might not project to the horizontal position of the trunk. A further source of confusion is that maps of geolocated stems tend to contain many more trees than can be seen from above, and the GNSS geolocation error of stems can vary among neighbouring trees. For all these reasons, it can be challenging to align geolocated stem measurements to a close-up photo of a canopy tree through spatial distance alone.

We recommend that the spatial co-registration between individual canopy trees and close-up photos is best made from above. In particular, we recommend segmenting individual crown boundaries from the RGB orthomosaic used to plan the close-up photo mission, or another one that is co-aligned. This can either be done manually (Araujo et al., 2020), or using machine learning models that can now perform well even in forests not represented in the training data (Baudchon et al., 2025). Such crown segmentations provide important context about tree crown diameter, from which diameter at breast height could also be estimated (Jucker et al., 2017, 2022). In addition, close-up photos can be acquired for these selected segmented trees to determine taxonomic identity. Close-up photos are easily linked to polygons of delineated crowns from the orthomosaic (which are perfectly spatially aligned with each other if using our workflow) via a simple spatial intersection operation. These sources of information (i.e. crown size, estimated stem diameter, and/or species) can provide additional sources of information about the tree to link it with its ground measurements, beyond the use of spatial coordinates alone. Once this is done, the high geolocation accuracy of both the close-up photos and the RGB orthomosaic, as well as the crown segmentation polygons that are now augmented with ground measurements, enables these data to be used as “canopy truthing” (as opposed to “ground truthing”) for other geospatial products.

### Drone safety considerations

We strongly recommend that the pilot becomes familiar with the theory and practice of RTK positioning before venturing into RTK mapping missions over tropical forest canopies, and especially close-up photo missions, which present the highest risk of drone collision with trees. The pilot should first perform RTK drone test flights in safe, open locations with no nearby obstacles. We recommend first completing mock mapping and close-up photo missions in a safe setting where the pilot has perfect visibility and control of the drone at all times to understand the drone behavior throughout the missions.

In our experience, the majority of issues that arise during close-up photo missions have to do with human errors related to RTK positioning of the drone, especially errors related to elevation. Because the drone acquires close-up photos at exactly 6 m above the canopy, the margin for error is small since the drone is very close to trees. Our recommendations are to: (1) check that the base station coordinates are the correct, expected ones before every flight, (2) wait for the drone to obtain RTK FIX before starting the close-up photo mission, (3) ensure that RTK signal is maintained during the entire mission, (4) ensure that the safe take-off and return-to-home (RTH) altitudes are set high enough (e.g., 100 m) to avoid a collision with any tree. In addition, the pilot should pay close attention to the camera view of the drone as it is approaching each tree for a close-up photo, especially the first few trees. Generally speaking, if the intended drone position for the first tree is correct, then everything should work well for all subsequent trees in the mission. The drone first approaches the tree at a faster downward speed up to 16 m above the tree, then will momentarily pause and make a slower approach to the tree up to 6 m. The pilot should pay attention to the downward distance to objects (the canopy) estimated by the drone built-in camera vision / obstacle avoidance system as the drone is moving close to the tree and be ready to press “Pause” and cancel the mission (RTH) if the drone seems to continue to move down beyond 6 m. When this happens, this is always due to the drone RTK status or base station elevation not having been correctly set using the steps listed above.

Therefore, we recommend that, depending on the region, each pilot—along with their co-pilot and field assistants—develop standardized pre- and post-flight checklists to minimize oversights and operational errors that could compromise flight safety. The list of instructions we send to our pilots is available in Appendix S1. Furthermore, we strongly advise that drone operators, particularly those working in dense tropical forests, conduct advanced training that includes simulations of potential in-flight emergencies (e.g., RTK signal loss, disconnection from the aircraft). Such preparation will ensure that the team has a clear and well-rehearsed contingency plan to safely recover the drone in the event of unexpected incidents.

Finally, an important safety consideration is to use the M3E obstacle avoidance system to prevent the drone from accidentally colliding with a tree. Given that **`harpia`** extracts the maximum elevation in a 3 m circular buffer around the selected location, we suggest using 3 m as the warning horizontal distance, and 2 m as the braking distance. For downward obstacle avoidance, we suggest using the maximum braking distance, which is 2 m. The upward distance is not critical since in theory there should never be any obstacles above the drone, but we still recommend turning it on and using a relatively large braking and warning distance (e.g., a few m) for additional safety.

### Past and future optimisations

Our first version of **`harpia`** was less efficient in that the drone would climb back up to the same predetermined safe altitude between all trees. This made it particularly inefficient in hilly locations. This first version was also limited in that the elevations for the close-up photos were calculated in reference to the take-off altitude, which limited us to this particular take-off site. By contrast, our newer, more efficient version uses ellipsoidal elevations directly so is no longer constrained to a particular take-off site, and follows a much more efficient flight path between trees. This second version is the one that is currently available at https://github.com/traitlab/harpia.

Further work could optimize flight paths between trees even further, to make it even more efficient. Other potential features could include options to acquire more than one close-up photo per selected tree crown polygon, including from different camera angles. In the future, we also plan to extend our workflow to the new DJI Matrice 4 Enterprise model (M4E), which has two major upgrades compared to the M3E: (1) the M4E has a 48 MP telephoto camera instead of the 12 MP pixel camera of the M3E, which should lead to even more visual detail to assist taxonomic identification, and (2) the new D-RTK3 GNSS base station has better signal transmission capacity and can relay the video signal if installed in a high position, which could help to extend the operational range of our workflow even further.

## Conclusions

In this study, we have presented a drone-based workflow to automatically acquire close-up, ultra-high resolution photos of tree crowns and lianas in forest canopies to support tropical botanical and ecological studies (https://youtu.be/80goMEifpc4). Our workflow is based around the DJI Mavic 3 Entreprise (M3E) drone, which is portable, relatively affordable and accessible to many organisations, as well as fairly easy to use. Our workflow allows one pilot to map ∼200 ha of forest in one day, and then acquire close-up photos from up to 300 trees per day on subsequent days. We have shown that these close-up photos allow expert botanists to reliably identify trees and lianas, often to species level. To program these close-up photos missions, we provide **`harpia`**, an open-source Python library (https://github.com/traitlab/harpia). We believe that our workflow could augment and accelerate tropical tree and liana surveys, and support phenological studies. Although our workflow was developed for tropical work, it could be extended to any other forest or vegetation type.

## Acknowledgements

We thank the administration and research staff from the field stations and sites where we conducted this research and developed our workflow: Tiputini Biodiversity Station (TBS), Barro Colorado Island (BCI), Adolfo Ducke Reserve, Tropical Silviculture Experimental Station (EEST), Station de biologie des Laurentides (SBL), AmazonFACE. In particular, we thank Gabriel Lanthier, Ciara Wirth, Carlos Valle, Carla Larrea, Esteban Rivera, Flavia Costa, and Juliana Schietti. We also thank members of the Limelight Rainforest team with whom we won the XPRIZE Rainforest competition for their camaraderie and support. In particular, we are grateful to Arthur Ouaknine, Mélisande Teng, Johanna Varner, Eric Fortune, Ryan Bixenman, and Outreach Robotics staff. This work was supported financially by a PRF3 grant “AI, Biodiversity, and Climate Change” from IVADO, by NSERC Discovery funds to E.L., by the Canada Research Chair in Plant Functional Biodiversity to E.L., and by a donation from the Population Biology Foundation. We also gratefully acknowledge financial support from the Simons Foundation award 429440 for the work in Panama and Universidad San Francisco de Quito for the financial and other support at the Tiputini Biodiversity Station. We thank Milton Garcia and César Gutierrez for assisting with drone permitting and data collection in Panama, and Salomón Aguilar and Ernesto Campos for assisting with species identification in Panama. We are also grateful to the Tiputini Biodiversity Station team for their support in arranging research permits in Ecuador, and to Niro Higuchi and INPA staff for arranging the research permit in Brazil. We thank the Ministry of the Environment of Ecuador for granting permit No. MAATE-ARSFC-2025-0319, and the Ministry of Science, Technology and Innovation of Brazil for granting permit CNPq No. 01300.003568/2024-46.

## Author contributions

Our study brings together authors from all countries where the research took place. Efforts were made to work collaboratively with local partners working at the sites. All authors were engaged early on with the research to ensure that the diverse sets of perspectives they represent were considered. Employment and authorship opportunities were offered to local scientists, technicians, and (para)botanists who contributed to the study. Local partners were provided with drone equipment and training to build research capacity in high-resolution remote sensing of forest canopies. E.L, A.C.G, V.L.F., and G.T. conceived the ideas, and designed the methodology. E.L., A.C.G., G.T., M.H., A.B., A.W., V.V., P.S., F.V., J.R., L.A.S., J.I., A.S., H.B. and J.G. collected drone imagery data and provided feedback on the data acquisition pipeline. J.V., F.A., A.H., D.B., J.C.I., G.R.T., and D.M.M. contributed to species identifications. H.M.L., G.R.T., T.R.W., D.M.M. and J.Q.C. supported data acquisition. A.C.G. and E.L. analysed the data. E.L. wrote the manuscript. All authors contributed critically to the drafts and gave final approval for publication.

## Conflict of interest

The authors have no conflict of interest to declare.

## Notes

https://github.com/traitlab/harpia

https://youtu.be/80goMEifpc4

### Competing Interest Statement

The authors have declared no competing interest.

### Summary of Updates

Some author names and affiliations were corrected.

