## Appendix S1 for "Seeing the forest and the trees: a workflow for automatic acquisition of ultra-high resolution drone photos of tropical forest canopies to support botanical and ecological studies"

### Appendix S1. Quick instructions for mapping and waypoint missions

1. Set up the DJI D-RTK 2 base, making sure it is correctly positioned on the reference point and level (check the 2 level bubbles).

2. Place the drone on a flat surface for takeoff and turn it on. Turn on the drone remote controller.

3. Ensure that the parameters of the mission to be carried out are correct.

   ### For mapping missions

   a. Real-time terrain monitoring mode should be activated for a height of 50 to 60 metres (usually).

   b. Choose a safe take-off altitude according to the topography of the terrain. This value should be at least similar to or higher than the mission altitude.

   c. Flight speed and course angle should be adjusted to minimise flight time. Select the highest possible speed according to the other parameters, and the best course angle to minimise mission time (flight lines should be parallel to the longest side of the mission polygon).

   d. In the advanced settings, the lateral overlap should be 75% (80% may be preferable in certain situations) and 85% for the frontal overlap. It may be preferable to add a margin of 10 to 20 metres if your polygon covers a specific area and you wish to have clear boundaries of this area when generating the orthomosaic, but the margin is not necessary in all cases. Select photo capture mode by distance.

   ### For waypoint missions (close-up photos)

   a. Open the mission after importing it and confirm that the mission corresponds to what is expected. Close the mission without saving it. The mission preview should stay blank if you do not save it.

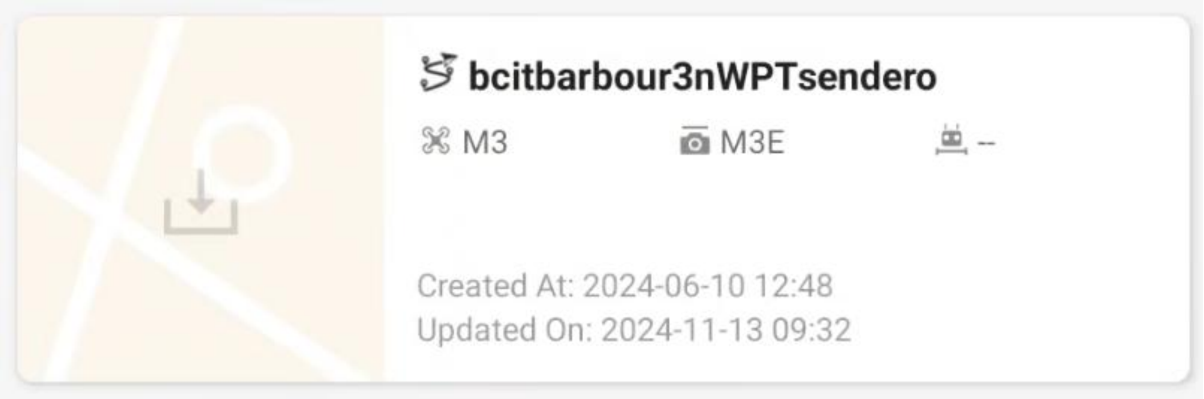


4. Open the camera view to check the drone's RTK parameters before takeoff. **Make sure the base position is correctly entered - it is very important!** You can modify the base position in the advanced settings by entering the code '123456'.

5. Open the mission to be carried out and launch the mission.

#### For mapping missions

a. Make sure the height of the RTH is sufficient for the topographical conditions.

b. Set the RTH point according to the drone's position before takeoff.

c. It is possible to deactivate the brakes on takeoff and landing, **but it is very important to reactivate them as soon as the drone is in the air.**

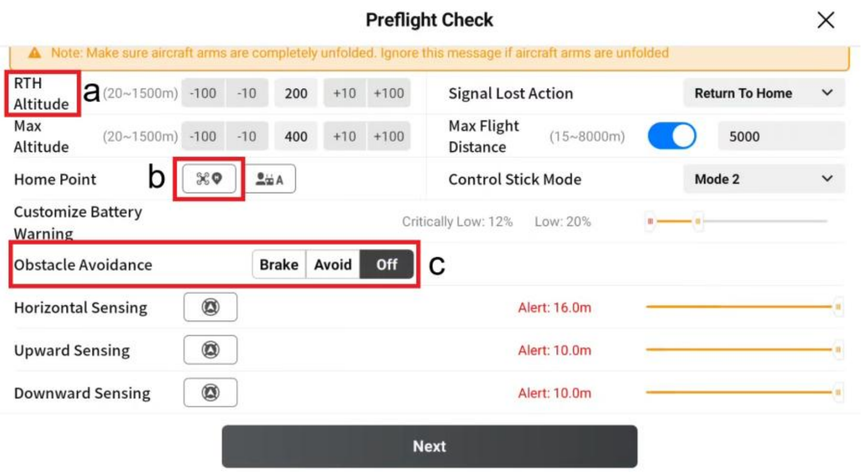

d. Ensure that the drone returns to the RTH point when the mission is completed, but continues the mission if the signal is lost.

e. Make sure the shutter speed is set to 1/1000 (for cloudy conditions) to 1/1250 (for sunny conditions).

f. Photo correction must be deactivated and white balance set to automatic mode.

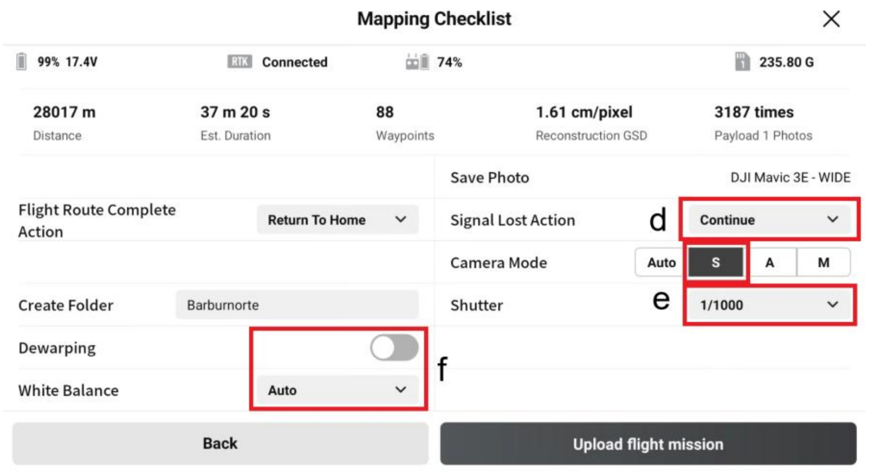


**g. Focus must be in manual mode (MF) during the flight.**

#### For waypoint missions (close-up photos)

a. Make sure the height of the RTH is sufficient for the topographical conditions.

b. Set the RTH point according to the drone's position before takeoff.

c. **It is possible to deactivate the brakes during take-off and landing, but it is very important to reactivate them as soon as the drone is in the air. The bottom brake (downward sensing, red line) is the most important, and should be set to brake at a distance of 2 meters.The lateral brake (horizontal sensing, red line) can block the drone if the value is too high, so it is preferable to keep the value at 2-3 meters. Alert threshold (yellow lines) can be kept at maximum value for all brakes.**

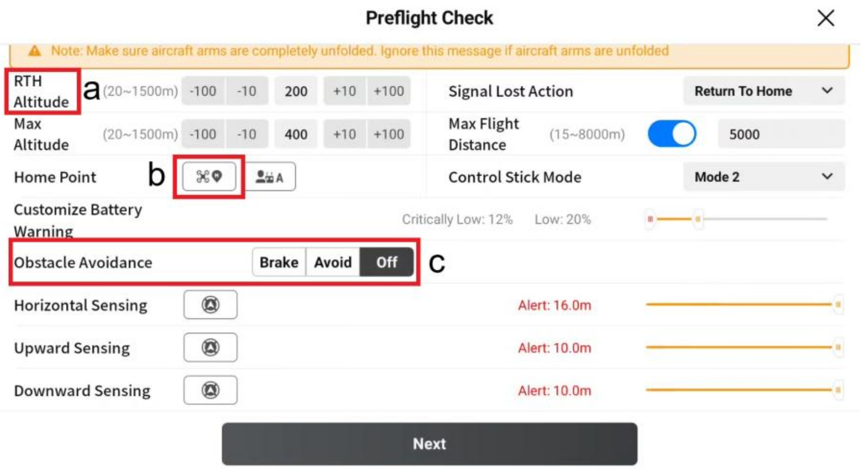


d. Ensure that the drone returns to the RTH point when the mission is completed, but continues the mission if the signal is lost.

**e. Make sure the shutter speed is set to 1/160.**

f. Set the white balance to automatic mode. In very cloudy conditions, if the photo becomes bluish, you can change the white balance to cloudy.

**g. Focus must be on continuous mode (AFC) during the flight.**

### A short list of the most important points:

#### For mapping missions

1. Make sure base position is correct
2. Make sure the shutter speed is set to 1/1000 (for cloudy conditions) to 1/1250 (for sunny conditions)
3. Focus must be in manual mode (MF) during the flight

#### For waypoint missions (close-up photos)

1. Make sure base position is correct
2. Make sure the shutter speed is set to 1/160
3. In very cloudy conditions, if the photos become bluish, you can change the white balance to cloudy
4. Focus must be on continuous mode (AFC) during the flight
